# Agreement between two large pan-cancer CRISPR-Cas9 gene dependency datasets

**DOI:** 10.1101/604447

**Authors:** Joshua M. Dempster, Clare Pacini, Sasha Pantel, Fiona M. Behan, Thomas Green, John Krill-Burger, Charlotte M. Beaver, Scott T. Younger, Victor Zhivich, Hanna Najgebauer, Felicity Allen, Emanuel Gonçalves, Rebecca Shepherd, John G. Doench, Kosuke Yusa, Francisca Vazquez, Leopold Parts, Jesse S. Boehm, Todd R. Golub, William C. Hahn, David E. Root, Mathew J. Garnett, Francesco Iorio, Aviad Tsherniak

## Abstract

Genome-scale CRISPR-Cas9 viability screens performed in cancer cell lines provide a systematic approach to identify cancer dependencies and new therapeutic targets. As multiple large-scale screens become available, a formal assessment of the reproducibility of these experiments becomes necessary. We analyzed data from recently published pan-cancer CRISPR-Cas9 screens performed at the Broad and Sanger institutes. Despite significant differences in experimental protocols and reagents, we found that the screen results are highly concordant across multiple metrics with both common and specific dependencies jointly identified across the two studies. Furthermore, robust biomarkers of gene dependency found in one dataset are recovered in the other. Through further analysis and replication experiments at each institute, we found that batch effects are driven principally by two key experimental parameters: the reagent library and the assay length. These results indicate that the Broad and Sanger CRISPR-Cas9 viability screens yield robust and reproducible findings.

## Introduction

Despite recent advances in cancer science, most cancer patients still have no clinical indications for approved targeted therapies^1^. Expanding precision oncology to the general patient population will require identifying and exploiting many new genomic targets. To tackle this problem, large-scale pharmacogenomic screenings have been performed across panels of human cancer cell lines^2,3^. The advent of genome editing by CRISPR-Cas9 technology has allowed extending these studies beyond currently druggable targets with precision and scale^4,5^. Pooled CRISPR-Cas9 screens employing genome-scale libraries of single guide RNAs (sgRNAs) are being performed on growing numbers of cancer *in vitro* models^6–12^. The output of these screens can be used to identify and prioritize new cancer therapeutic targets^13^. However, fully characterizing genetic vulnerabilities in cancers is estimated to require thousands of genome-scale screens^14^.

We present a comparative analysis of datasets derived from the two largest independent CRISPR-Cas9 based gene-dependency screening studies in cancer cell lines published to date^13,15,16^, part of the *Cancer Dependency Map* effort^17,18^. The aim of this analysis was to assess the concordance of these datasets and that of the analytical outcomes they yield when investigated individually. To this aim, we designed a computational strategy including comparisons at different levels of data-processing and abstraction: from gene-level dependencies to molecular markers of dependencies, and genome-scale cell line profiles of dependencies. Lastly, we shed light on the differences in the experimental settings that give rise to batch effects across independent studies of this kind, discerning between biological and technical confounding factors.

## Results

### Overview of datasets and comparison strategy

We compared two sets of pooled genome-scale CRISPR-Cas9 drop out screens in cancer cell lines, generated at the Broad Institute and the Sanger Institute through experimental pipelines (detailed in **Fig. 1a, Supplementary Table 1** and **Supplementary text**), considering 147 cell lines and 16,733 genes screened independently by both institutes (**Supplementary Table 2**). We performed comparisons of individual gene dependency scores, quantifying the reduction of cell viability upon gene inactivation via CRISPR-Cas9 targeting; of profiles of such scores across cell lines (gene dependency profiles); as well as of profiles of such scores across genes in individual cell lines (cell line dependency profiles).

**Figure 1:**
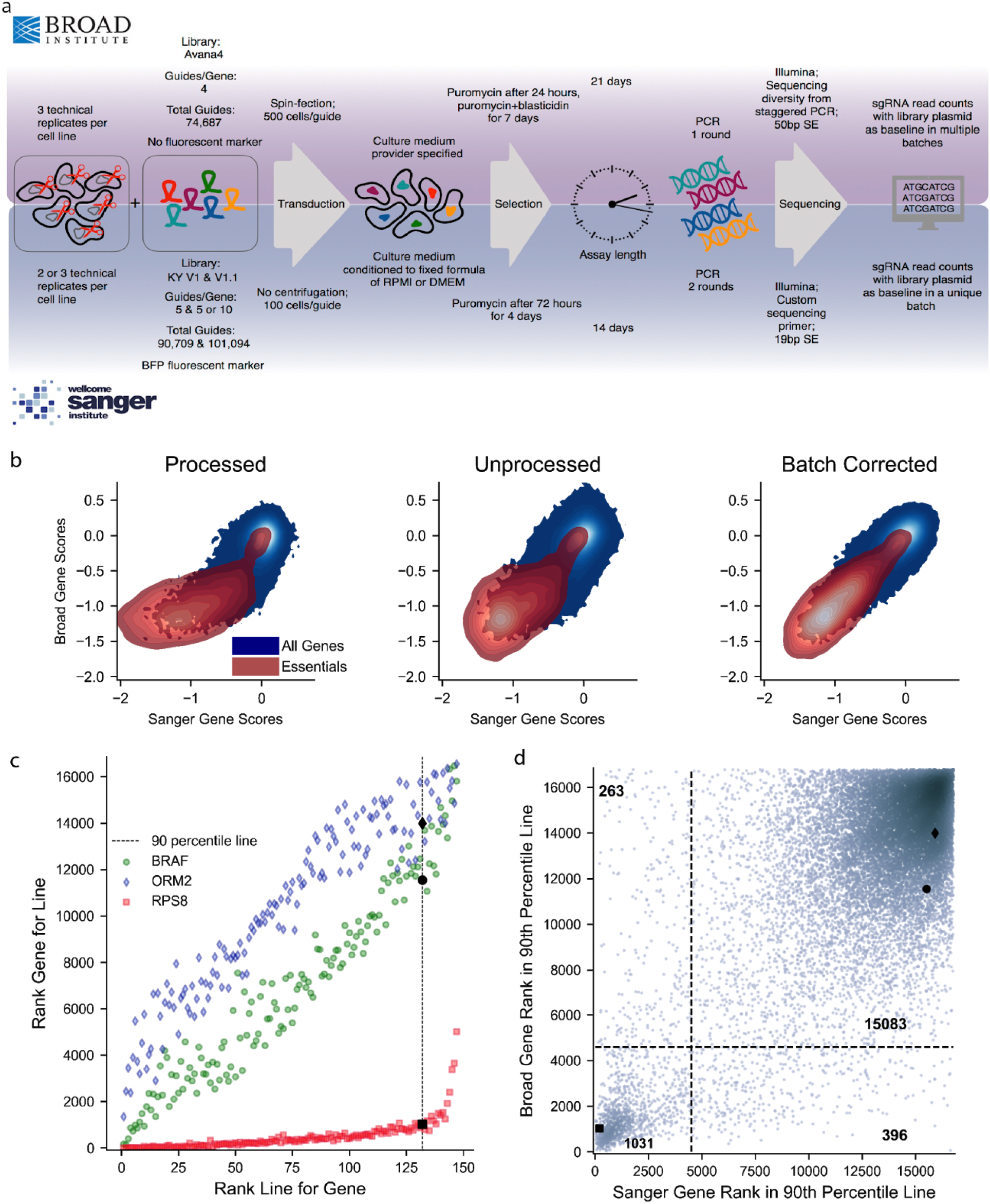
Comparison of experimental protocols and gene score results, (**a**) Comparison of experimental settings and reagents used in the experimental pipelines underlying the two compared datasets, (**b**) Densities of individual gene scores in the Broad and Sanger datasets, across processing levels. The distributions scores for previously identified essential genes^20^ are shown in red. (**c**) Examples of the relationship between a gene’s score rank in a cell line and the cell line’s rank for that gene using Broad unprocessed gene scores, with gene ranks in their 90th percentile least dependent lines highlighted. For the common dependency RPS8, even its 90th percentile least dependent cell line still ranks the gene among the strongest of its dependencies, (**d**) Distribution of gene ranks for the 90th-percentile of least dependent cell lines for each gene in both datasets. Black dotted lines indicate natural thresholds at the minimum gene density along each axis.

We calculated gene dependency scores using three different strategies. First, we considered fully processed gene scores, available for download from the Broad^17^ and Sanger^13,18^ Cancer Dependency Map web-portals^17,13,18^ (*processed* data). Because data processing pipelines vary significantly between the two datasets, we also examined minimally processed gene scores, generated by computing median sgRNA abundance fold changes for each gene *(unprocessed* data). Lastly, we applied an established empirical Bayesian batch correction method (ComBat)^19^ to the unprocessed gene scores to remove experimental batch effects between the datasets. ComBat aligns gene means and variances between the datasets, thereby eliminating simple batch effects. We refer to this form of the data as the *batch-corrected* gene scores.

### Agreement of gene dependency scores

We found concordant gene scores across all genes and cell lines with Pearson correlation = 0.658, 0.627, and 0.765, respectively for processed, unprocessed and batch-corrected data (*p* values below machine precision in all cases, *N =* 2,465,631, **Fig. 1b**). The reproducibility of gene scores between the two datasets can be considered a function of two variables: the mean dependency across all cell lines for each gene (relevant to infer common dependencies), and the patterns of scores across cell lines for each gene (relevant to predict selective oncology therapeutic targets). Mean gene scores among all cell lines showed excellent agreement (**Supplementary Fig. 1a**), with Pearson correlation = 0.784, 0.818, and 0.9997 respectively for processed, unprocessed and batch-corrected data (p below machine precision in all cases; *N =* 16,773).

We further tested whether it was possible to recover consistent sets of common dependencies. To this end, we defined as “common dependencies” those genes that rank among the top dependencies when considering only their 90th percentile of least dependent cell lines, with the score threshold for “top” dependencies determined by the local minimum in the data (**Fig. 1c**). For the unprocessed data, the Broad and Sanger jointly identify 1,031 common dependency genes (**Supplementary Table 3**). 260 putative common dependencies were only identified by the Sanger and 397 were only identified by the Broad (Cohen’s kappa = 0.737, Fisher’s exact test p below machine precision, *N =* 16,773, **Fig. 1d**).

### Agreement of selective gene dependency profiles across cell lines

In both studies, most genes show little variation in their dependency scores across cell lines. Thus we expect low shared variance even if most scores are numerically similar between the datasets^21^. Accordingly, we focused on a group of genes for which the score variance across lines is of potential biological interest. These are genes whose dependency profile suggests a strong biological selectivity in at least one of the two unprocessed datasets, identified using the NormLRT test introduced in McDonald *et al*^22^. We call these 49 genes *Strongly Selective Dependencies* (SSDs) (**Supplementary Table 4**). We evaluated the agreement between gene score patterns using Pearson’s correlations to test the reproducibility of selective viability phenotypes. **Fig. 2a** illustrates the score patterns for the example cancer genes MDM4 (*R* = 0.820, *p* = 6.91 x 10^-37^), KRAS (*R* = 0.765, *p* = 1.66 x 10^-29^), CTNNB1 (*R* = 0.803, *p* = 1.92 x 10^-34^), and SMARCA4 (*R* = 0.664, *p* = 4.61 x 10^-20^) with unprocessed data (*N* = 147). For SSDs and unprocessed data, the median correlation was 0.633 and 84% of SSDs showed a correlation greater than 0.4. Five SSDs showed a correlation below 0.2 (ABHD2, CDC62, HIF1A, HSPA5, C17orf64), which are discussed further below. As expected, correlation across datasets for all genes was lower (median *R* = 0.187, 8.34% genes with *R* > 0.4).

**Figure 2:**
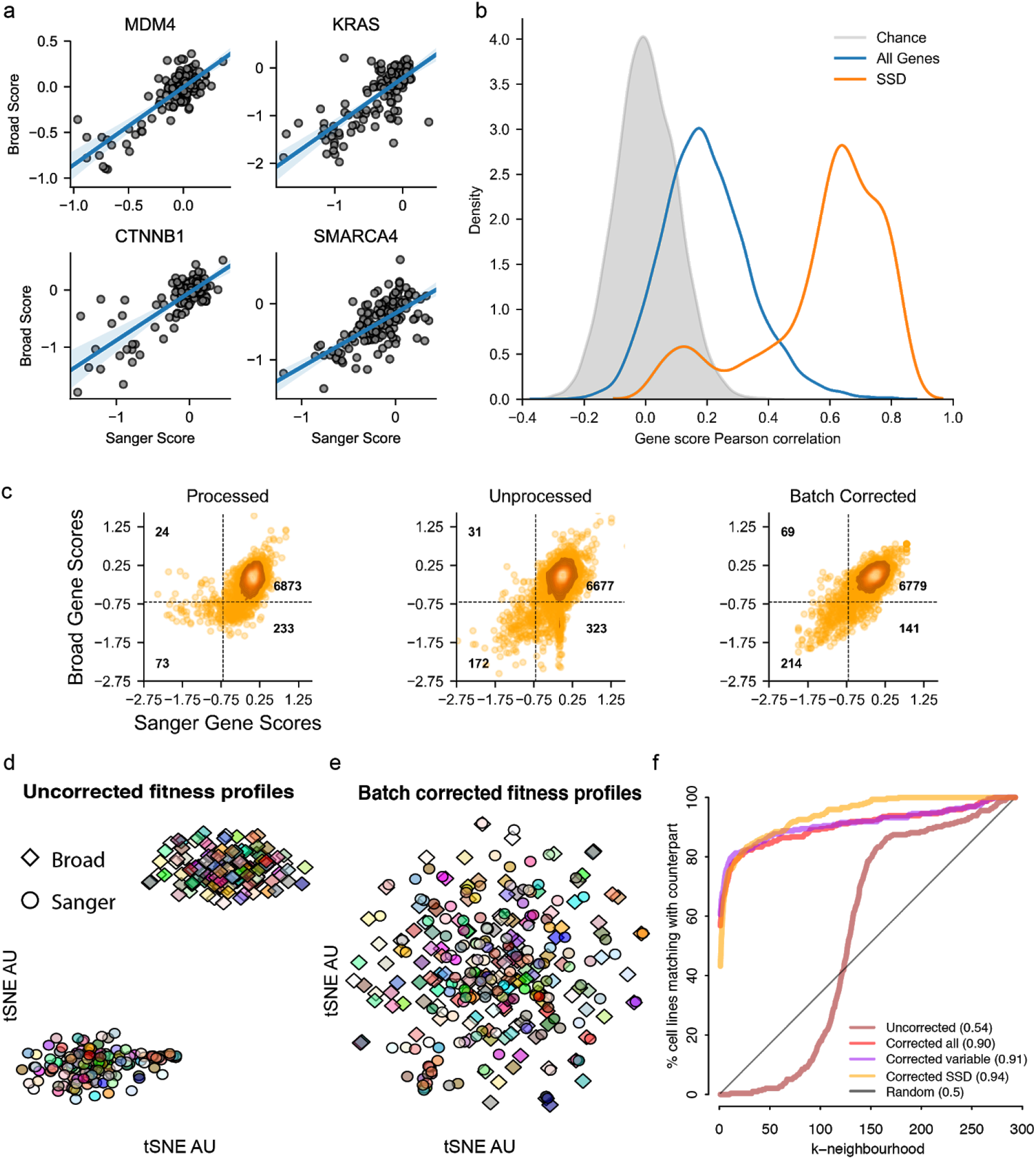
Reproducibility of gene and cell line dependency profiles, (**a**) Score pattern examples for selected known cancer genes, (**b**) Distribution of correlations of scores for individual genes in unprocessed data, (**c**) Gene dependency scores for Strongly Selective Dependencies across all cell lines, with the threshold for calling a line dependent set at −0.7. (**d**) t-SNE clustering of cell lines in unprocessed data using the correlation between gene scores. Colors represent the cell line while shape denotes the dataset of origin, (**e**) The same as in (d) but for data batch-corrected using ComBat. (**f**) Recovery of a cell line’s counterpart in the other dataset before (Uncorrected) and after correction (Corrected). The k-nearest neighborhood shows the percentage of cell lines whose matching counterpart in the other data set is within its k-nearest cell lines. Similarity between all cell line pairs is defined by the Pearson correlation and nAUC values are shown in brackets. We show the recovery using three different gene sets to calculate the correlation between cell lines. First, using all genes (Uncorrected and Corrected all), second using genes that are dependencies for at least one cell line (corrected variable) and third using the Strongly Selective Dependencies (SSD) genes.

One important use of these screens is to consistently classify cells as dependent or not dependent on selective dependencies. Therefore, we evaluated the agreement of the Broad and Sanger datasets on identifying cell lines that are dependent on each SSD gene. We classified cell lines as dependent on a given gene in the case of a dependency score < −0.7. Genes with larger scores are dominated by a single large group at zero (**Fig. 2c**). The area under the receiver-operator characteristic (AUROC) for recovering binary Sanger dependency on SSDs using Broad dependency scores was 0.940 in processed data, 0.963 in unprocessed data, and 0.965 in corrected data; to recover Broad binary dependency from Sanger scores, AUROC scores were = 0.933, 0.859, and 0.967 respectively. The recall of Sanger-identified dependent cell lines in Broad data was 0.700 with precision equal to 0.252 for processed data, 0.847 and 0.347 for unprocessed data, and 0.756 and 0.603 for batch-corrected data (**Supplementary Fig. 1b**).

Agreement is higher than could be expected by chance under all processing regimes (Fisher’s exact p = 4.16 x 10^-41^ in processed, 1.37 x 10^-86^ in unprocessed, and 2.96 x 10^-215^ in batch-corrected data; *N* = 7,203). A large proportion of Broad-exclusive dependent cell lines (56.2 % in processed data and 42.7% in unprocessed data) were due to the single gene HSPA5, which is an SSD in Sanger data but a common dependency in Broad data. Examining SSDs individually, we found median Cohen’s kappa for sensitivity to individual SSDs of 0.437 in processed, 0.661 in unprocessed, and 0.735 in batch-corrected data. In unprocessed data, 65% of SSDs had Cohen’s kappa greater than 0.4, as opposed to 0.17% seen by chance (**Supplementary Fig. 1c**).

### Agreement of cell line dependency profiles

Previous literature on reproducibility highlighted the importance of considering agreement along both the perturbation and cell line axes of the data^23–25^. We assembled a combined dataset of cell line dependency profiles from both studies and computed all possible pairwise correlation distances between them, using genes that were dependencies in at least one cell line *(variable* genes). A t-distributed stochastic neighbor embedding (tSNE)^26^ visualization derived from these distance scores is shown in **Fig. 2d**. For the uncorrected data, we observed a perfect clustering of the dependency profiles by their study of origin, confirming a major batch effect. However, following batch correction, we observed integration between studies and increased proximity of cell lines from one study to their counterparts in the other study (**Fig. 2e**). To quantify agreement, for each cell line dependency profile in one dataset, we ranked all the others (from both datasets) based on their correlation distance to the profile under consideration. For batch-corrected data, 175 of 294 (60%) cell lines dependency profile from one study have their counterpart in the other study as the closest (first) neighbor, and 209 of 294 (71%) cell lines having it among the five closest neighbors (area under the normalized Recall curve - nAUC - averaged across all profiles = 0.91 for batch-corrected data, and = 0.54 for uncorrected data, **Fig. 2f**). Similar results were obtained across dependency profiles restricted to different sets of genes, with the best performance for the SSD genes (nAUC=0.94) and the worst for all genes (nUAC=0.90). The percentage of cell lines matching closest to their counterparts in the other study was 57% for the variable gene set and 43% for SSD genes. Further, the tSNE plots for each gene set showed similar improvement after correction (**Supplementary Fig. 2a-b**).

The batch correction also aligned numbers of significant (at 5% FDR) dependencies across cell lines between the two datasets (median number of dependencies 2,109 and 1,717 before, and 2,053 and 1,950 after correction, for Broad and Sanger respectively, **Supplementary Fig. 3a**). The average proportion of dependencies detected in both studies over those detected in at least one study also increased across cell lines from 47.75% to 59.14%. Furthermore, the correlation between cell lines after correction rose above the correlation within each individual screen for each gene set considered (**Supplementary Fig 3b**). We finally examined whether the residual disagreement in corrected data might be related to screen quality. We assessed screen quality by computing true positive rates (TPRs) for recovering common essential genes in each cell line with a fixed 5% false discovery rate (FDR), determined from the distribution of nonessential genes in the cell line. We found that mean screen quality is a strong predictor of screen agreement for both the uncorrected and batch-corrected data sets (p-values 2.06 x 10^-35^, 4.74 x 10^-35^ and adjusted R-squared 0.65, 0.64 for uncorrected and batch-corrected respectively; **Supplementary Fig. 3c**).

### Agreement of gene dependency biomarkers

A selective dependency is of limited therapeutic value unless it can be reliably associated with an informative molecular feature of cancer (*biomarker*). Following a similar approach to that presented in^21^, we performed a systematic test for molecular-feature/dependency associations on the two datasets. Cell lines were split into two groups based on the status of 587 molecular features derived from Iorio *et al.^27^*, encompassing somatic mutations in high-confidence cancer driver genes, amplifications/deletions of chromosomal segments recurrently altered in cancer, hypermethylated gene promoters, microsatellite instability status and the tissue of origin of the cell lines (**Supplementary Table 5**). For each feature in turn, all SSD genes were sequentially *t*-tested for significant differences in dependency scores between the obtained two groups of cell lines.

These tests yielded 71 out of 29,350possible significant associations (FDR < 5%, ΔFC < −1) between molecular features and gene dependency when using the Broad unprocessed data, and 90 when using the Sanger unprocessed data (**Supplementary Table 6**). Of these, 55 (77% of the Broad associations and 61% of the Sanger ones) were found in both datasets (FET p-value = 9.08 x10^-133^, **Fig. 3a** and **Supplementary Tables 6-7**). The concordance between the associations identified by each study was proportional to the threshold used to define significance. This was assessed by considering for each study, in turn, the associations in a fixed quantile of significance and measuring the tendency of these associations to be among the most significant in the other study (**Fig. 3b**). Further, the overall correlation between differences in gene depletion FCs was equal to 0.763, and 99.2% of associations had the same sign of differential dependency across the two studies.

**Figure 3:**
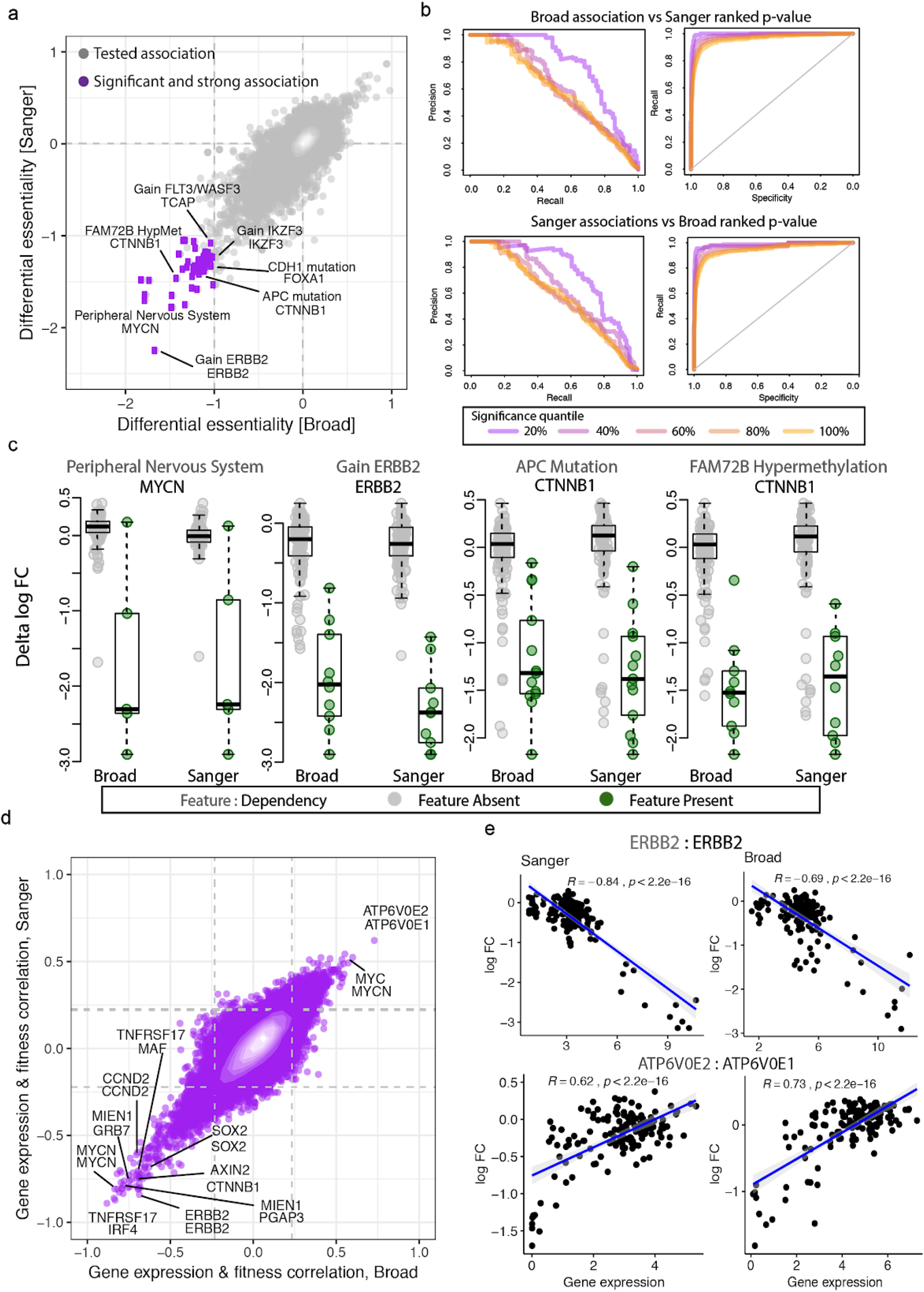
Reproducibility of biomarkers. (**a**) Results from a systematic association test between molecular features and differential gene dependencies (of the SSD genes) across the two studies. Each point represents a pair consisting of a molecular feature (on the first line) and a gene dependency (second line.) (**b**) Precision/Recall and Recall/Specificity curves obtained when considering as true positives the associations falling in a fixed top quantile of significance in one of the studies and a classifier based on the p-values of the associations from the other study, (**c**) Examples of significant statistical associations between genomic features and differential gene dependencies across the two studies, (**d**) Results from a systematic correlation test between gene expression and dependency of SSD genes across the two studies. Labelled points show the gene expression marker on the first line and gene dependency on the second line, (**e**) Examples of significant correlations between gene expression and dependencies in both studies

Gene dependency associations identified with both datasets included expected as well as potentially novel hits. Examples of expected associations included increased dependency on ERBB2 in ERBB2-amplified cell lines, and increased dependency on beta-catenin in APC mutant cell lines. A potentially novel association between FAM72B promoter hypermethylation and beta-catenin was also identified (**Fig. 3c**).

We also considered gene expression to mine for biomarkers of gene dependency using RNA-seq datasets maintained at Broad and Sanger institutes. To this aim, we considered as potential biomarkers 1,987 genes from intersecting the top 2,000 most variable gene expression levels measured by either institute. Clustering the RNA-seq profiles revealed that each cell line’ transcriptome matched closest to its counterpart from the other institute (**Supplementary Fig. 4a**).

We correlated the gene expression level for the most variably expressed genes to the gene dependency profiles of the SSD genes. Systematic tests of each correlation showed significant associations between gene expression and dependency (**Fig. 3d**). As with the genomic biomarkers, we found a strong overall correlation between gene expression markers and SSD genes dependency across datasets, Pearson’s correlation 0.804 and significantly high overlap between gene expression biomarkers identified in each dataset (Fisher’s exact test p-value below machine precision). We observed both positive and negative correlations; for example, ERBB2 dependency score was positively correlated with its expression, while ATP6V0E1 showed significant dependency when its paralog ATP6V0E2 had a low expression (**Fig. 3e**).

### Elucidating sources of disagreement between the two datasets

Despite the concordance observed between the Broad and Sanger datasets, we found batch effects in the unprocessed data both in individual genes and across cell lines. Although the bulk of these effects are mitigated by applying an established correction procedure^28^, their cause is an important experimental question. We enumerated the experimental differences between datasets (**Fig. 1a**) to identify likely causes of batch effects. The choice of sgRNA can significantly influence the observed phenotype in CRISPR-Cas9 experiments, implicating the differing sgRNA libraries as a likely source of batch effect^29^. Additionally, previous studies have shown that some gene inactivations results in cellular fitness reduction only in lengthy experiments^11^. Accordingly, we selected the sgRNA library and the timepoint of viability readout for primary investigation as causes of major batch effects across the two compared studies.

To elucidate the role of the sgRNA library, we examined the data at the level of individual sgRNA scores. The correlation between log fold change patterns of reagents targeting the same gene (“co-targeting”) across studies was related to the selectivity of the average dependency of that gene (as quantified by a “Likelihood Ratio Test” – normLRT – score^22^, **Fig. 4a):** a reminder that most co-targeting reagents show low correlation because they target genes exerting little phenotypic variation. However, even among SSDs there was a clear relationship between sgRNA correlations within and between datasets (*p* = **4.9** x 10^-10^, *N =* **49; Fig. 4b**). In such cases, poor reagent efficacy at one or both datasets may explain the discrepancy. We estimated the efficacy of each sgRNA in both libraries using Azimuth 2.0^29^ which uses only information about the genome in the region targeted by the sgRNA. We found that among genes identified as common dependencies in either dataset, mean sgRNA depletion indeed had a strong relationship to its Azimuth estimated efficacy (**Fig. 4c**). When we examined SSDs, we found that reagent efficacy likely explains some differences, for example in EIF3F (common essential in Sanger screens, nonscoring in Broad screens) and MDM2 (strongly selective in Broad screens, correlated but not strongly selective in Sanger screens) (**Fig. 4d**).

**Figure 4:**
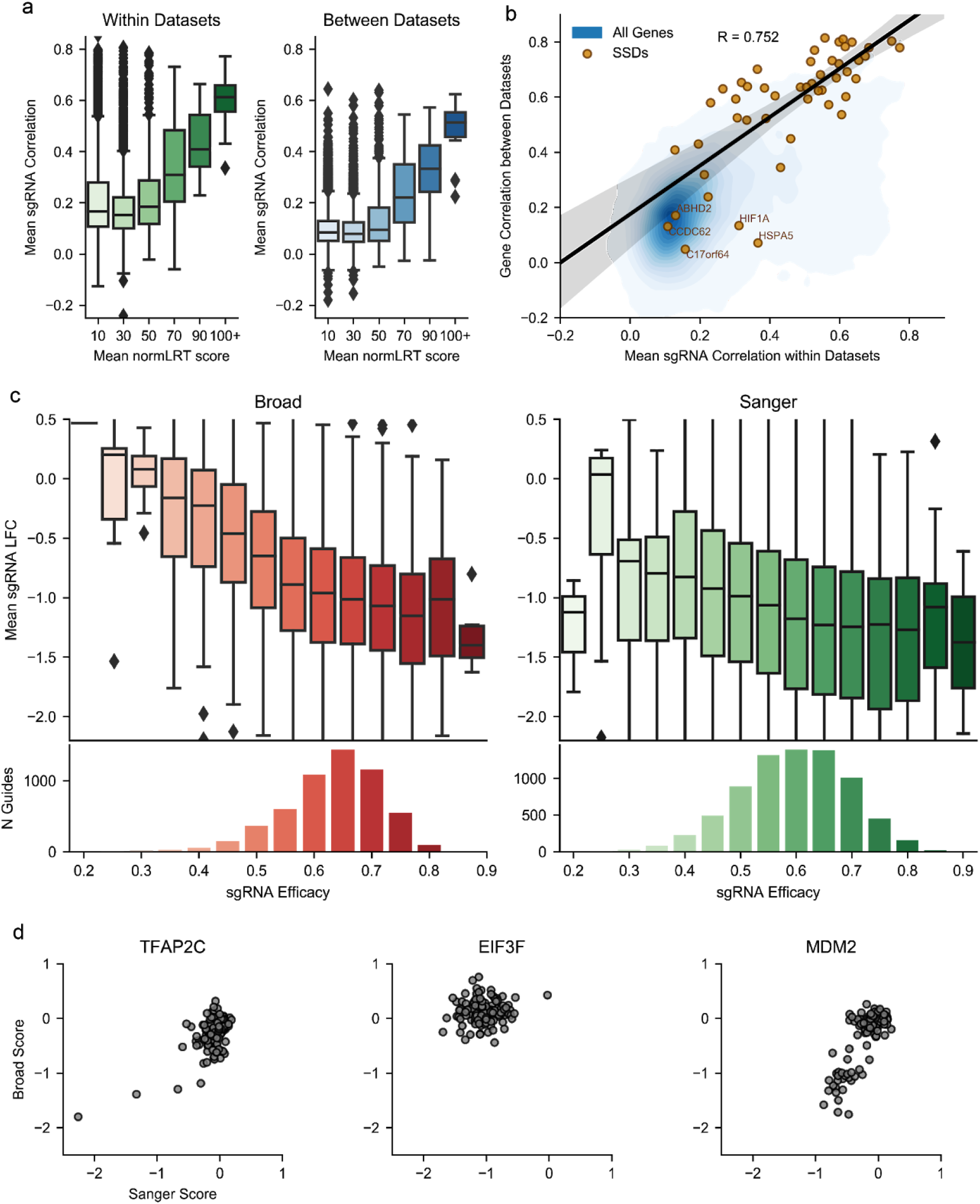
Influence of reagent library on measured gene dependency, (**a**) Distributions of sgRNA correlation for sgRNAs targeting genes with varying NormLRT scores within each dataset and between them. Each gene is binned according to the mean of its NormLRT score across the two datasets. The y axis reports the average of all correlations between pairs of sgRNAs that belong to the same dataset and target that gene, (**b**) Relationship between sgRNA correlation within datasets and gene correlation between datasets. The linear trend is shown for SSD genes, (**c**) The mean depletion of guides targeting common dependencies across all replicates vs Azimuth estimates of guide efficacy, (**d**) Comparison of Broad and Sanger unprocessed gene scores for genes matching (1) SSD with highest minimum MESE across both libraries, (2) common dependency in either dataset and greatest difference between KY and Avana MESE, (3) SSD with worst KY MESE.

We next investigated the role of different experimental timepoints on the screens’ agreement. Given that the Broad used a longer assay length (21 days versus 14 days) we expected differences to be observed between late dependencies across the datasets. Therefore, we compared the distribution of gene scores for genes known to exert a loss of viability effect upon inactivation at an early- or late-time *(early or late dependencies*)^11^. While early dependencies have similar score distributions in both datasets (median average score −0.781 at the Sanger and −0.830 at the Broad), late dependencies are more depleted at the Broad with median average score −0.402 compared to −0.269 for the Sanger screens (**Fig. 5a**). Unlike differences in sgRNA efficacy, timepoint effects are expected to lead to uniformly greater signal (typically depletion) in the Broad data and to be related to the biological role of late dependencies. We functionally characterized, using gene ontology (GO), genes that were exclusively detected as depleted in individual cell lines (at 5% FDR), in one of the two studies, excluding genes with significantly different sgRNA efficacies between libraries. Results showed 29 gene ontology categories significantly enriched in the Broad-exclusive dependencies genes (Broad-exclusive GO terms) for more than 50% of cell lines (**Fig. 5b** and **Supplementary Table 8**). The Broad-exclusive enriched GO terms included classes related to mitochondrial and RNA processing gene categories and other gene categories previously characterized as late dependencies^11^. In contrast, no GO terms were significantly enriched in the Sanger-exclusive common dependencies in more than 30% of cell lines.

**Figure 5:**
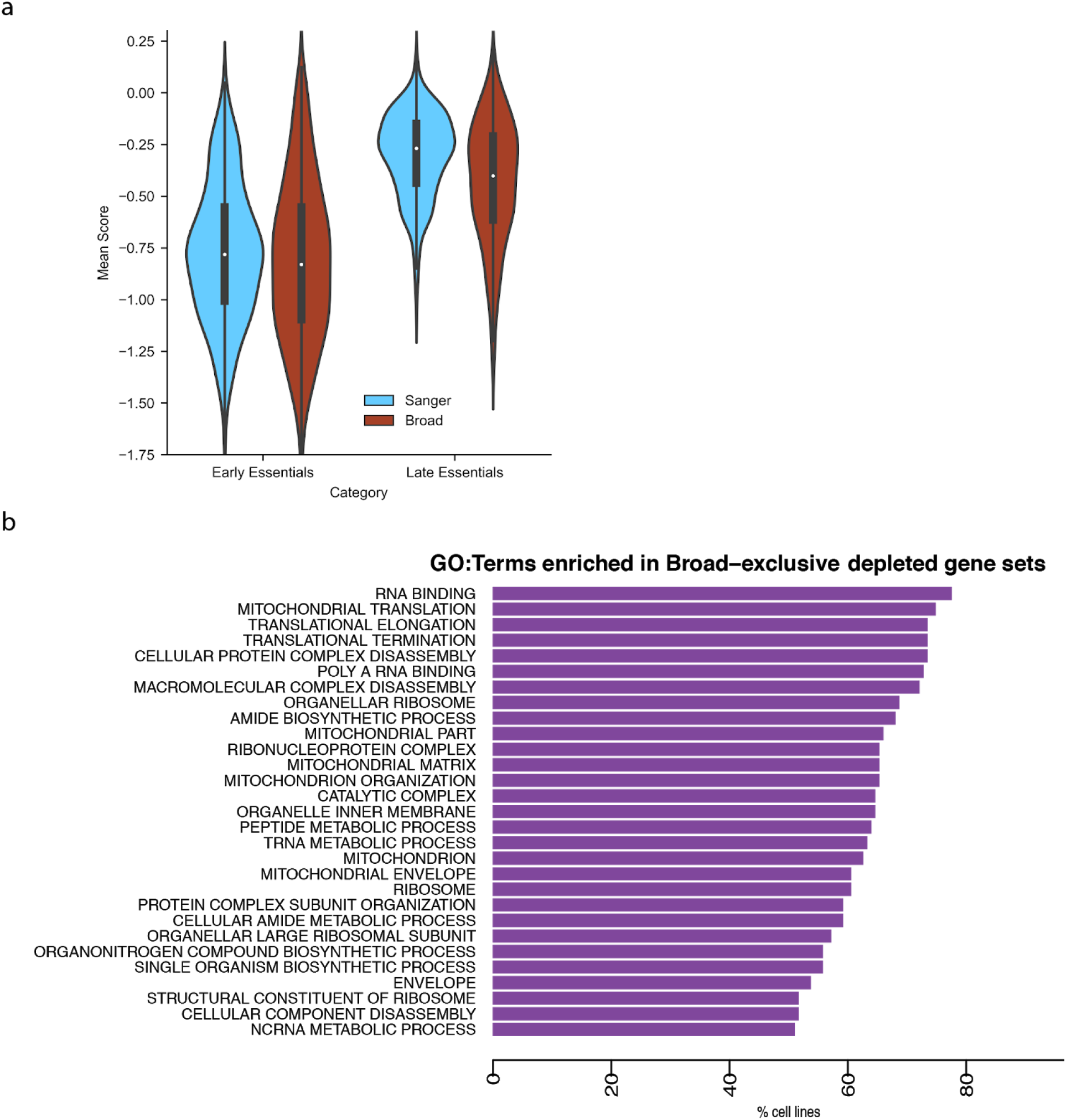
Influence of Time point, (**a**) Distribution of early and late common dependency scores in the Broad and Sanger datasets averaged across cell lines, (**b**) GO enrichment annotations of Broad-exclusive common dependencies not accounted for by estimated sgRNA efficacy.

### Batch effect sources: Experimental Verification

To verify that batch effects between the datasets can be removed by changing library and the readout time-point, we undertook replication experiments independently at Broad and Sanger institutes, where these factors were systematically permuted. The Broad sequenced cells collected from its original HT-29 and JIMT-1 screens at the 14-day timepoint and conducted an additional screen of these cell lines using the KY1.1 library with readouts at days 14 and 21. The Sanger used both the Broad’s and the Sanger’s clones of HT-29 to conduct a new KY screen and an Avana screen with readouts at days 14 and 21. Principal component analysis (PCA) of the concatenated unprocessed gene scores, including replication screens, showed a clear institute batch effect dominating the first principal component. By highlighting replication screens, we found that this effect is principally due to library choice, with time-point playing a smaller role (**Fig. 6a, Supplementary Fig. 5a**). Changing from Sanger to Broad clones of HT-29 had minimal impact. We examined the change in gene score profile for each screen caused by changing either library or time-point while keeping other conditions constant. Gene score changes induced by either library or timepoint alterations were consistent across multiple conditions (**Fig. 6b**). Sanger-exclusive common dependencies were strongly enriched for genes that became more depleted with the KY library, and Broad-exclusive common dependencies were enriched among genes more depleted with the Avana library (**Supplementary Fig. 5b**). Late dependencies were strongly enriched among genes that became more depleted in the later time-points, while early dependencies were not (**Supplementary Fig. 5c**). We compared the deviations in gene score between Broad and Sanger screens under different conditions, first comparing Broad original and replication screens of HT-29 (Fig. 6c) and JIMT-1 (**Supplementary Fig. 5d**) to the original Sanger screens of the same cell line. Matching Sanger’s library and time-point reduces the variance of gene scores in HT-29 from 0.0486 to 0.0252 and in JIMT-1 from 0.0556 to 0. 0260. Specifically, matching library and time-point removed most of the average gene score change (batch effect) between institutes, as indicated by the low correlation of the remaining gene score differences in the replication screens with the average gene score change. We next compared Sanger original and replication screens of HT-29 to the Broad original HT-29 screen. Matching library and time-point successfully detrended the data in this case as well; however, the Sanger Avana screens of HT-29 contained considerable excess noise, causing these screens to have higher overall variance from the Broad than the original screens (0.0486 vs 0.115). Nonetheless, the replication experiments confirm that the majority of batch effects between datasets are driven by library and time-point.

**Figure 6:**
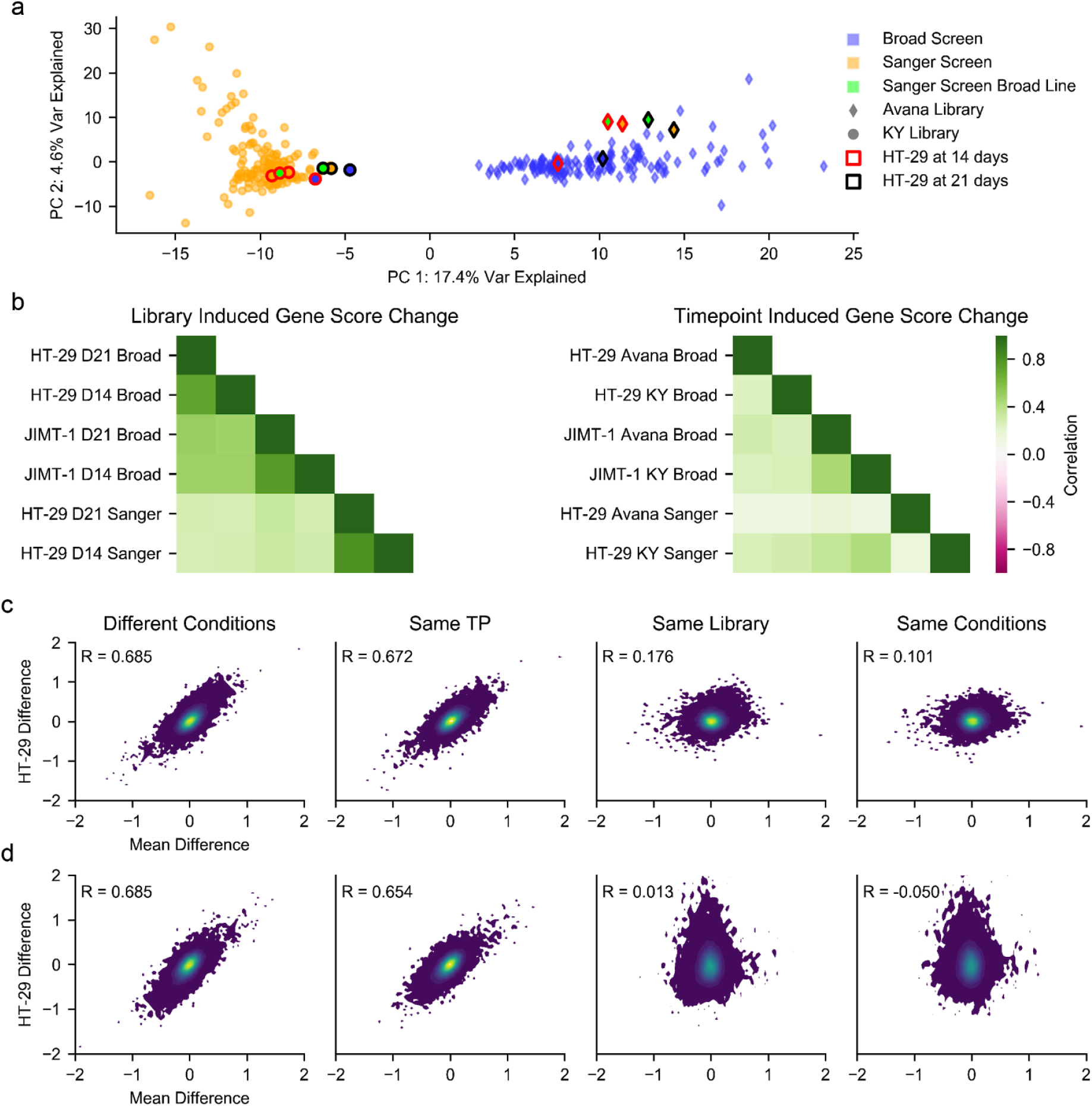
Results of replication experiments, (**a**) original and replication screens from each institute plotted by their first two principal components. HT-29 screens are highlighted. Axes are scaled to the variance explained by each component, (**b**) Correlations of the changes in gene score caused when changing a single experimental condition, (**c**) The difference in unprocessed gene scores between Broad screens of HT-29 and the original Sanger screen (Sanger minus Broad), beginning with the Broad’s original screen and ending with the Broad’s screen using the KY library at the 14-day timepoint. Each point is a gene. The horizontal axis is the mean difference of the gene’s score between the Sanger and Broad original unprocessed datasets, (**d**) A similar plot taking the Broad’s original screen as the fixed reference and varying the Sanger experimental conditions (Broad minus Sanger).

## Discussion

Providing sufficient experimental data to adequately sample the diversity of human cancers requires high-throughput screens. However, the benefits of large datasets can only be exploited if the underlying experiments are reliable and robustly reproducible. In this work, we survey the agreement between two large, independent CRISPR-Cas9 knock-out datasets, generated at the Broad and Sanger institutes.

Our findings illustrate a high degree of consistency in estimating gene dependencies between studies at multiple levels of data processing, albeit with the longer duration of the Broad screens leading to stronger dependencies for a number of genes. The datasets are concordant in identifying common dependencies and identifying mean dependency signals. Their agreement is also striking in the more challenging task of identifying which cell lines are dependent on selective dependencies. Indeed, when we compared the two datasets at the level of gene dependency markers we found consistent results at the level of common informative molecular features, as well as with respect to their quantitative strength.

We observed that a source of observed disagreement across the compared dataset is due to diffuse batch effects visible when the whole profiles of individual cell lines are compared. Such effects can be readily corrected with standard methods without compromising data quality, thus making possible integration and future joint analyses of the two compared datasets. Furthermore, much of this batch effect can be decomposed into a combination of two experimental choices: the sgRNA library and the duration of the screen. The effect of each choice on the mean depletion of genes is readily explicable and reproducible, as shown by screens of two lines performed at the Broad using the Sanger’s library and screen duration. Consequently, identifying high-efficacy reagents and choosing the appropriate screen duration should be given high priority when designing CRISPR-Cas9 knock-out experiments.

## Methods

### Collection and Preprocessing of Data

#### “Unprocessed” Gene Scores

Read counts for the Broad were taken from avana_public_19Q1^30^ and filtered so that they contained only replicates corresponding to overlapping cell lines and only sgRNAs with one exact match to a gene. Read counts for Sanger were taken from Behan *et al.^13^* and similarly filtered, then both read counts were filtered to contain only sgRNAs matching genes common to all versions of the data. In both cases, reads per million (RPM) was calculated and an additional pseudo-count of 1 added to the RPM. Log fold change was calculated from the reference pDNA. In the case of the Broad, both pDNA and screen results fall into distinct batches, corresponding to evolving PCR strategies. Cell lines sequenced with a given batch were matched to pDNA profiles belonging to the same batch. Multiple pDNA RPM profiles in each batch were median-collapsed to form a single profile of pDNA reads for each batch. Initial gene scores for each replicate were calculated from the median of the sgRNAs targeting that replicate. Each replicates initial gene scores for both Broad and Sanger were then shifted and scaled so the median of nonessential genes in each replicate was 0 and the median of essential genes in each replicate is negative one^20^. Replicates were then median-collapsed to produce gene-by cell-line matrices.

#### “Processed” Gene Scores

Broad gene scores are taken from avana_public_19Q1 gene_effect^30^ and reflect CERES^31^ processing. The scores were filtered for genes and cell lines shared between institutes and with the unprocessed data, then shifted and scaled so the median of nonessential genes in each cell line was 0 and the median of essential genes in each cell line was −1^20^. Sanger gene scores were taken from the quantile-normalized averaged log fold-change scores and globally rescaled by a single factor so that the median of essential genes across *all* cell lines is negative one.^20^

#### “Batch-Corrected” Gene Scores

The unprocessed sgRNA log FCs were mean collapsed by gene and replicates. Data were quantile normalized for each institute separately before processing with ComBat using the R package sva. One batch factor was used in ComBat defined by the institute of origin. The ComBat corrected data was then quantile normalized to give the final batch-corrected data set.

#### Alternate Conditions

Screens with alternate libraries, cell lines, and timepoints were processed similarly to the “Unprocessed” data above.

#### Gene Expression Data

Gene expression log_2_(Transcript per million+1) data was downloaded for the Broad from Figshare for the Broad data set. For the Sanger dataset, we used reads per kilobase million (RPKM) expression data from the iRAP pipeline. We added a pseudo-count of 1 to the RPKM values and transformed to log_2_. Gene expression values are quantile normalized for each institute separately. For the Sanger data, Ensembl gene ids were converted to Hugo gene symbols using BiomaRt package in R.

### Guide Efficacy Estimates

On-target guide efficacies for the single-target sgRNAs in each library were estimated using Azimuth 2.0^29^ against GRCh38.

### Comparison of All Gene Scores

Gene scores from the chosen processing method for both Broad and Sanger were raveled and Pearson correlations calculated between the two datasets. 100,000 gene-cell line pairs were chosen at random and density-plotted against each other using a Gaussian kernel with the width determined by Scott’s rule^32^. All gene scores for essential genes were similarly plotted in **Fig. 1b**.

### Comparison of Gene Means

Cell line scores for each gene in both Broad and Sanger datasets with the chosen processing method were collapsed to the mean score, and a Pearson correlation calculated.

### Gene Ranking, Common Essential Identification

For each gene in the chosen dataset, its score rank among all gene scores in its 90th percentile least depleted cell line was calculated. We call this the gene’s 90th percentile ranking. The density of 90th-percentile rankings was then estimated using a Gaussian kernel with width 0.1 and the central point of minimum density identified. Genes whose 90th-percentile rankings fell below the point of minimum density were classified as essential.

### Identification of Selective Gene Sets

Selective dependency distributions across cell lines are identified using a “Likelihood Ratio Test” as described in *McDonald et al^22^*. For each gene, the log-likelihood of the fit to a normal distribution and a skew-t distribution is computed using R packages MASS^33^ and sn^34^, respectively. In the event that the default fit to the skew-t distribution fails, a two-step fitting process is invoked. This involves keeping the degrees of freedom parameter (***v***) fixed during an initial fit and then using the parameter estimates as starting values for a second fit without any fixed values. This process repeats up to 9 times using ***v*** values in the list (2, 5, 10, 25, 50, 100, 250, 500, 1000) sequentially until a solution is reached. The numerical optimization methods used for the estimates do not guarantee the maximum of the objective function is reached. The reported LRT score is calculated as follows:

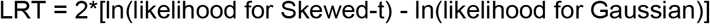

Genes with NormLRT scores greater than 100 and mean gene score greater than −0.5 in at least one institute’s unprocessed dataset were classified as SSDs. The cancer gene set was taken directly from Vogelstein *et al.^35^*

### Binarized Agreement of SSDs

SSD gene scores in both Broad and Sanger datasets with the chosen processing method were binarized at −0.7, with scores falling below this threshold indicating the sensitivity of the cell line on the chosen gene. Cohen’s kappa was calculated for each gene individually. Fisher’s exact test, precision, recall, and AUROC scores were calculated globally for all SSD sensitivities in the three data versions.

### Cell line agreement Analysis

To obtain the two dimensional visualisations of the combined dataset before and after batch correction and considering different gene sets, we computed the sample-wise correlation distance matrix and used this as input into the t-statistic Stochastic Neighbor Embedding (t-SNE) procedure^26^, using the *tsne* function of the tsne R package, with 1,000 iterations, a perplexity of 100 and other parameters set to their default value.

To evaluate genome-wide cell line agreement we considered a simple nearest-neighbor classifier that, for each dependency profile of a given cell line in one of the two studies, predicts its matching counterpart in the other study. This prediction was based on the correlation distance between one profile and all the other profiles. To estimate the performance of this classifier, we computed a Recall curve for each of the 294 dependency profiles in the tested dataset. Each of these curves was assembled by concatenating the number of observed true-positives amongst the first *k* neighbors of the corresponding dependency profile (for *k =* 1 to 293). We then averaged the 294 resulting Recall curves into a single curve and converted it to percentages by multiplying by 100/294. Finally we computed the area under the resulting curve and normalized it by dividing by 293. We considered the area under this curve (nAUC) as a performance indicator of the k-nn.

For the comparison of cell line profiles agreement in relation to initial data quality. First, to estimate the initial data quality we calculated True Positive Rates (TPRs, or Recalls) for the sets of significant dependency genes detected across cell lines, within the two studies. To this aim, we used as positive control a reference set of a priori known essential genes^12^. We assessed the resulting TPRs for variation before/after batch correction, and for correlations with the inter-study agreement.

### Biomarker Analysis

We used binary event matrices based on mutation data, copy number alterations, tissue of origin and MSI status. The resulting set of 587 features were present in at least 3 different cell lines and fewer than 144. We performed a systematic two-sample unpaired Student’s t-test (with the assumption of equal variance between compared populations) to assess the differential essentiality of each of the SSD genes across a dichotomy of cell lines defined by the status (present/absent) of each CFE in turn. From these tests we obtained p-values against the null hypothesis that the two compared populations had an equal mean, with the alternative hypothesis indicating an association between the tested CFE/gene-dependency pair. P-values were corrected for multiple hypothesis testing using Benjamini-Hochberg. We also estimated the effect size of each tested association by means of Cohen’s Delta, i.e. difference in population means divided by their pooled standard deviations. For gene expression analysis we calculated the Pearson correlation across the cell lines between the SSD gene dependency profiles and the gene expression profiles from each institute. Significance of the correlation was assessed using the t-distribution (n-2 degrees of freedom) and p-values multiple hypothesis corrected using the q-value method.

For the agreement assessment via ROC indicators (Recall, Precision and Specificity), for each of the two studies in turn we picked the most significant 20, 40, 60, 80 and 100% associations as true controls and evaluated the performance of a rank classifier based on the corresponding significance p-values obtained in the other study.

For the analysis involving transcriptional data, we used the RNA-seq data from each institute for overlapping cell lines, which includes some sequencing files that have been used by both institutes and processed separately.

### Rank-based dependency significance and agreement quantification

To identify significantly depleted genes for a given cell line, we ranked all the genes in the corresponding essentiality profiles based on their depletion logFCs (averaged across targeting guides), in increasing order. We used this ranked list to classify genes from two sets of prior known essential (*E*) and non-essential (*N*) genes, respectively^12^.

For each rank position *k*, we determined a set of predicted genes 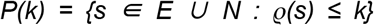, with 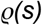 indicating the rank position of *s*, and the corresponding precision *PPV(k)* as:

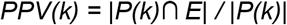

Subsequently, we determined the largest rank position *k*\* with *P*(*k*\*) ≥ 0.95 (equivalent to a False Discovery Rate (FDR) ≤ 0.05). Finally, a 5% FDR logFCs threshold *F\** was determined as the logFCs of the gene s such that 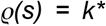, and we considered all the genes with a logFC < *F*\* as significantly depleted at 5% FDR level. For each cell line, we determined two sets of significantly depleted genes (at 5% FDR): *B* and *S*, for the two compared datasets, respectively. We then quantified their agreement using the Jaccard index^36^ *J*(*B,S*) = |*B* ∩ *S* |/| *B* ∪ *S*|, and defined their disagreement as *1* – *J*(*B,S*). Summary agreement/disagreement scores were derived by averaging the agreement/disagreement across all cell lines.

### sgRNA Correlations

Broad and Sanger log fold-changes for their original screens were median-collapsed to guide by cell line matrices. For each gene present in the unprocessed gene scores, a correlation matrix between all the sgRNAs targeting that gene in each guide by cell line matrix was computed. The mean of the values in this matrix for each institute, excluding the correlations of sgRNAs with themselves, was retained. The mean sgRNA correlation within institutes was then calculated from the mean of the Broad and Sanger sgRNA correlation matrix means. The mean sgRNA correlation between institutes for each gene was calculated from the mean of all possible pairs of sgRNAs targeting that gene with one sgRNA chosen from Sanger and one from Broad.

### Relating sgRNA Depletion and Efficacy

We chose the set of genes found to be essential in at least one unprocessed dataset. The log fold-change of guides targeting those genes in each dataset was calculated and compared to the guide’s estimated on-target efficacy.

### Timepoint Gene Ontology Analysis

We tested for enrichment of GO terms associated with genes showing a significant depletion in only one institute. To rule out the differences due to library, genes with significantly different guide efficacies were filtered from the analysis. Using the Azimuth scores average (mean) efficacy scores for each gene at each institute were calculated. A null distribution of differences in gene efficacy was estimated using genes not present in either institute specific sets (which were defined as depleted in at least 25% of cell lines). Institute specific genes greater than 2 standard deviations from the mean of the null distribution were removed.

For the filtered gene set prior known essential and non-essential gene sets from ^37^ were used to find significant depletions for each cell line and institute at 5% FDR. For each cell line, the genes identified as significantly depleted in only Broad or only Sanger were functionally characterized using Gene Ontology (GO) enrichment analysis^38^. To this aim, we downloaded a collection of gene sets (one for each GO category) from the Molecular Signature Database (MsigDB)^39^, and performed a systematic hypergeometric test to quantify the over-representation of each GO category for each set of study-exclusive dependency genes, per cell line. We corrected the resulting p-values for all the tests performed within each study using the Benjamini-Hochberg procedure^40^, and considered a GO category enriched in a cell line if the corrected p-value resulting from the corresponding test was < 0.05.

### Principal Component Analysis of the Batch Effect and Alternate Conditions

The Broad and Sanger unprocessed gene scores and the gene scores for the alternate conditions tested by both institutes were concatenated into a single matrix with a column for each screen. Principal components were found for the transpose of this matrix, where each row is a screen and each column a pseudogene. Components 1 and 2 were plotted for all original screens and the alternate screens for either HT-29 (**Fig. 6a**) or JIMT-1 (**Supplementary Fig. 6a**). The aspect ratio for the plot was set to match the relative variance explained by the first two principal components.

### Consistency of Timepoint and Library Effects on Gene Scores

To evaluate library differences, we took all screens that had been duplicated in each library with all other conditions (timepoint, clone, and screen location) kept constant. For each of these screens, we subtracted the gene scores of the version performed with the KY library from the version performed with the Avana library to create library difference profiles. For the case of Sanger’s day-14 KY screen of the Sanger HT-29 clone, two versions exist, the original and an alternative that was eventually grown out to 21 days. We used the alternate version of this screen to be consistent with the day 21 results. A correlation matrix of library difference profiles was then calculated and is plotted in the left of **Fig. 6b**. The procedure was repeated for timepoint differences, creating timepoint difference profiles by subtracting day 14 results from day 21 results for pairs of screen readouts that differed in timepoint but not library, clone, or screen location.

### Mitigating Differences in Gene Scores by Matching Experimental Conditions

For the cell line HT-29, we took Sanger’s original screen as a baseline. We then subtracted from this baseline from four Broad HT-29 screens: the original (Avana library at day 21), then with the Avana library at day 14, the KY library at day 21, and the KY library at day 14, generating four arrays indexed by gene which form the y-axes in the succession of plots in **Fig. 6c**. We also computed the mean score of each gene across all original Broad screens and subtracted it from the mean score of each gene across all the original Sanger screens to form the x-axis of all four plots. For each condition, the standard deviation of the HT-29 screen differences (y-axes) was computed along with the correlation of the HT-29 screen differences with the mean differences (x-axis). The plots themselves are Gaussian kernel density estimates. We repeated this process for JIMT-1 (**Supplementary Fig. 6d**) and then for HT-29 while swapping the roles of Broad and Sanger (**Fig. 6d**). For the Sanger alternate condition screens we used the Sanger clone of HT-29, and for its day 14 KY screen we used the Sanger’s original HT-29 screen.

### Replication Experiments

The replication screens at Broad and Sanger were performed using the normal current protocol of the respective institution^31^ except with respect to the specifically noted changes to library (and the associated primer sequences required for post-screen amplification of the sgRNA barcodes) and the timepoint.

## Data Availability

The data used for this manuscript have been posted to Figshare^41^.

## Acknowledgments

This work was funded by Open Targets (OTAR2-055) to F.I. and (OTAR015) to M.G. and K.Y., by the Wellcome Trust grant no. 206194 to M.G., by Wellcome and the Estonian Research Council (IUT 34-4) to L.P., by grants U01 CA176058 and U01 CA199253 to W.C.H and by the HL Snyder Foundation (W.C.H.).

