## Supplemental Data 1 for "Agreement between two large pan-cancer CRISPR-Cas9 gene dependency datasets"

### Supplementary material for “Agreement between two large pan-cancer CRISPR-Cas9 gene dependency datasets”

### Supplementary Tables

***Supplementary Table 1 - Experimental pipelines' specifications***

***Supplementary Table 2 - Cell lines screened in both studies***

***Supplementary Table 3 - Common dependencies in each dataset***

***Supplementary Table 4 - Gene NormLRT scores in both studies and genes used in the systematic statistical inference of essentiality markers***

***Supplementary Table 5 - Cell line molecular features used in the systematic inference of gene essentiality markers***

***Supplementary Table 6 - Dependency markers test results from the two individual studies***

***Supplementary Table 7 - Marker consistency via ROC analysis***

***Supplementary Table 8 - GO:terms enriched in the Broad-exclusive dependencies***

### Supplementary Data

**Supplementary Data can be accessed on Figshare**[1]**.**

### Supplementary Text

#### Overview of the experimental pipelines underlying the compared datasets

In both the Broad and Sanger studies, Cas9 expressing cell lines were generated using lentiviral delivery of a Cas9 expression vector containing a blasticidin resistance selection marker. Different minimal Cas9 activity levels were required at each site (45% at Broad; 75% at Sanger) for a cell line to be included in the genome scale CRISPR-Cas9 screening pipeline. Cell lines successfully exhibiting Cas9 activity above the respective thresholds were transduced with a pooled genome-scale sgRNA library in different numbers of technical replicates (3 replicates at Broad; 2 or 3 replicates at Sanger). In addition, in the Broad pipeline, for some cell lines, experimental replicates were performed transducing the library in different batches.[2]

The Broad screening pipeline utilised the Avana v4 library[2]; whereas the Sanger screening pipeline used one of two libraries, KY human v1.0[3] or KY human v1.1[4]. KY human v1.1 contains all sgRNAs included in KY human v1.0 plus an additional 1,000 non-targeting sgRNAs and 5 extra sgRNAs targeting a subset of ~500 genes from v1.0. There are noteworthy differences between Avana v4 and KY human libraries: length of sgRNAs; number of sgRNAs targeting each gene (Avana v4 has 4; KY human v1.0 has 5 & v1.1 has 5 or 10); total number of sgRNAs in the library (Avana v4 has 74,687; KY human v1.0 has 90,709 & v1.1 has 101,094).

Cell lines were transduced using different methods and aimed for different levels of library coverage at each site (the Broad pipeline used centrifugation aiming for transduction of 500 cells per sgRNA i.e. 500x coverage of the library; the Sanger pipeline used polybrene to facilitate transduction of 100 cells per sgRNA i.e. 500x coverage of the library). Antibiotic selection strategies for transduced cells also differed between the two sites (Broad added puromycin 24 hours post transduction followed by addition of concurrent blasticidin selection for 7 days; Sanger puromycin selection commenced 72 hours post transduction with 4 day duration). The overall duration of the assay also differed between the two sites (21 days at Broad; 14 days at Sanger). Genomic DNA was extracted using a similar method, followed by 2 unique sgRNA amplification and sequencing strategies. Broad pipeline employed a 1 step PCR method using multiple staggered PCR primer pairs to introduce diversity of amplicons to facilitate accurate sequencing. Sequencing tags were simultaneously added to each sample during this PCR step. Illumina 50bp single end sequencing was completed on each sample with resulting reads trimmed to include only the sgRNA sequence. Sanger pipeline employed a 2 step PCR strategy. Step 1 amplified the sgRNA using a single primer pair; step 2 added unique sequencing tags to each sample. Illumina 19bp single sequencing was completed using a custom primer that directs sequencing of the sgRNA alone.

The sgRNA read-counts resulting from the two pipelines (for each cell line replicate, together with the library plasmid read-counts), were used to assemble sgRNA-level and gene-level depletion log fold-change (logFCs) datasets as detailed in the Methods (one dataset per study). These were used as the starting point of our comparative analysis.

### Supplementary Figures

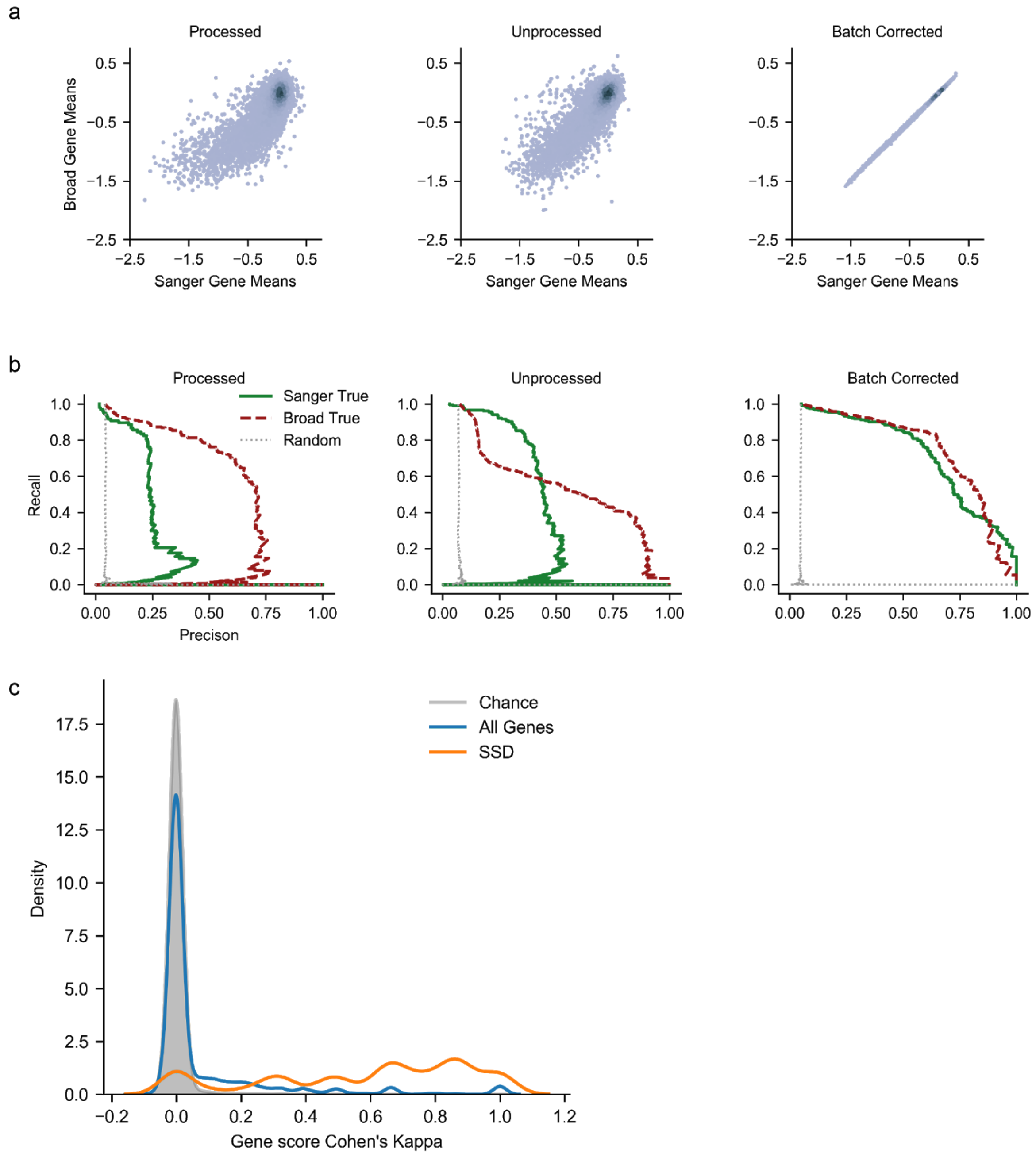


**Supplementary Figure 1: Gene Mean Dependency and Gene Selectivity**. **(a)** Comparison of gene mean scores between Broad and Sanger institutes under three different processing regimes. **(b)** Recall-precision curves for the sensitivity status of cell lines to SSD knockout, recovering Sanger statuses from Broad data and Broad statuses from Sanger. **(c)** Distribution of Cohen's kappa for different gene groups between unprocessed datasets.

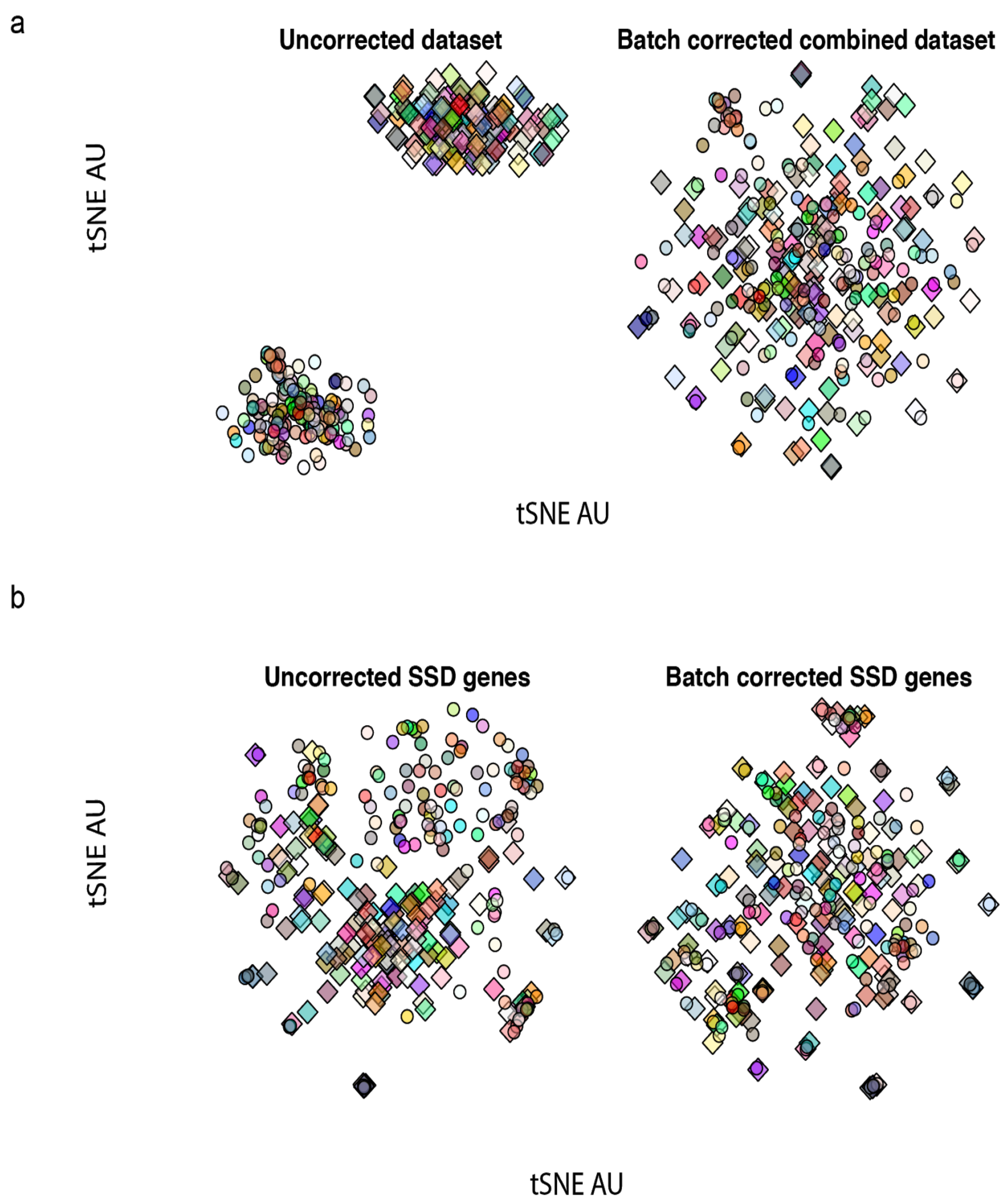


**Supplementary Figure 2**: **Batch correction assessment for different gene sets. (a)** tSNE plot for all genes, uncorrected and batch-corrected. **(b)** Same as (a) but using the Strongly Selective Dependency (SSD) genes.

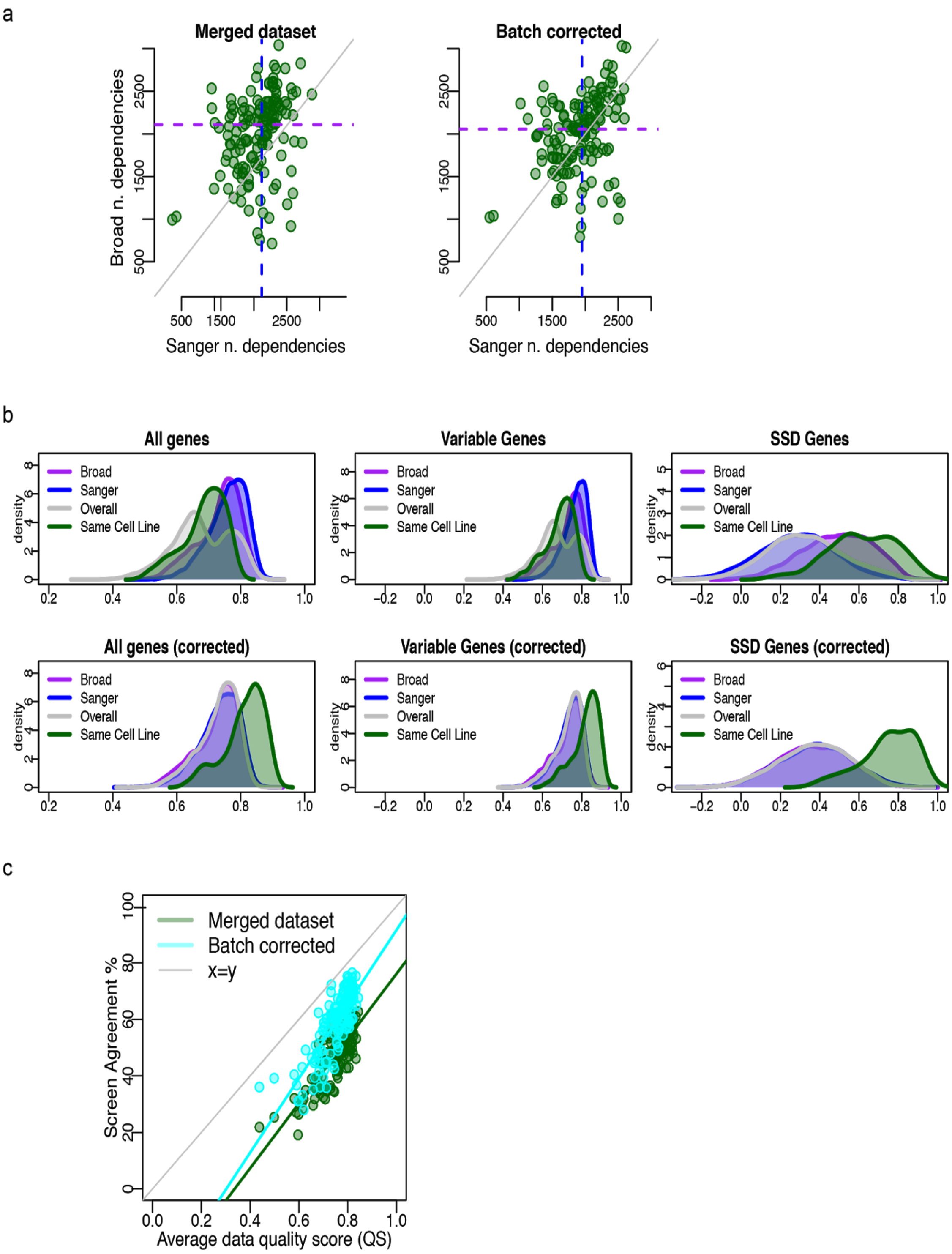


**Supplementary Figure 3**: **Reproducibility of gene and cell line profiles**. **(a)** Number of dependencies identified in each of the institutes for each cell line pair before and after correction. After correction the number of dependencies identified for each institute is in greater agreement. **(b)** Correlation scores between cell lines, overall for all cell line pairs, for all pairs of cell lines within institutes and between the same cell lines across institutes. This is shown for uncorrected data and batch-corrected data sets based on different gene sets.

Correlation between the same cell lines across studies is improved in all cases following correction with ComBat. **(c)** Relationship between average screen quality for a cell line pair and the agreement between them. For a cell line where the screen quality from both institutes is high the agreement of their dependency profiles is better than where one or both screens for a cell line is lower quality.

a

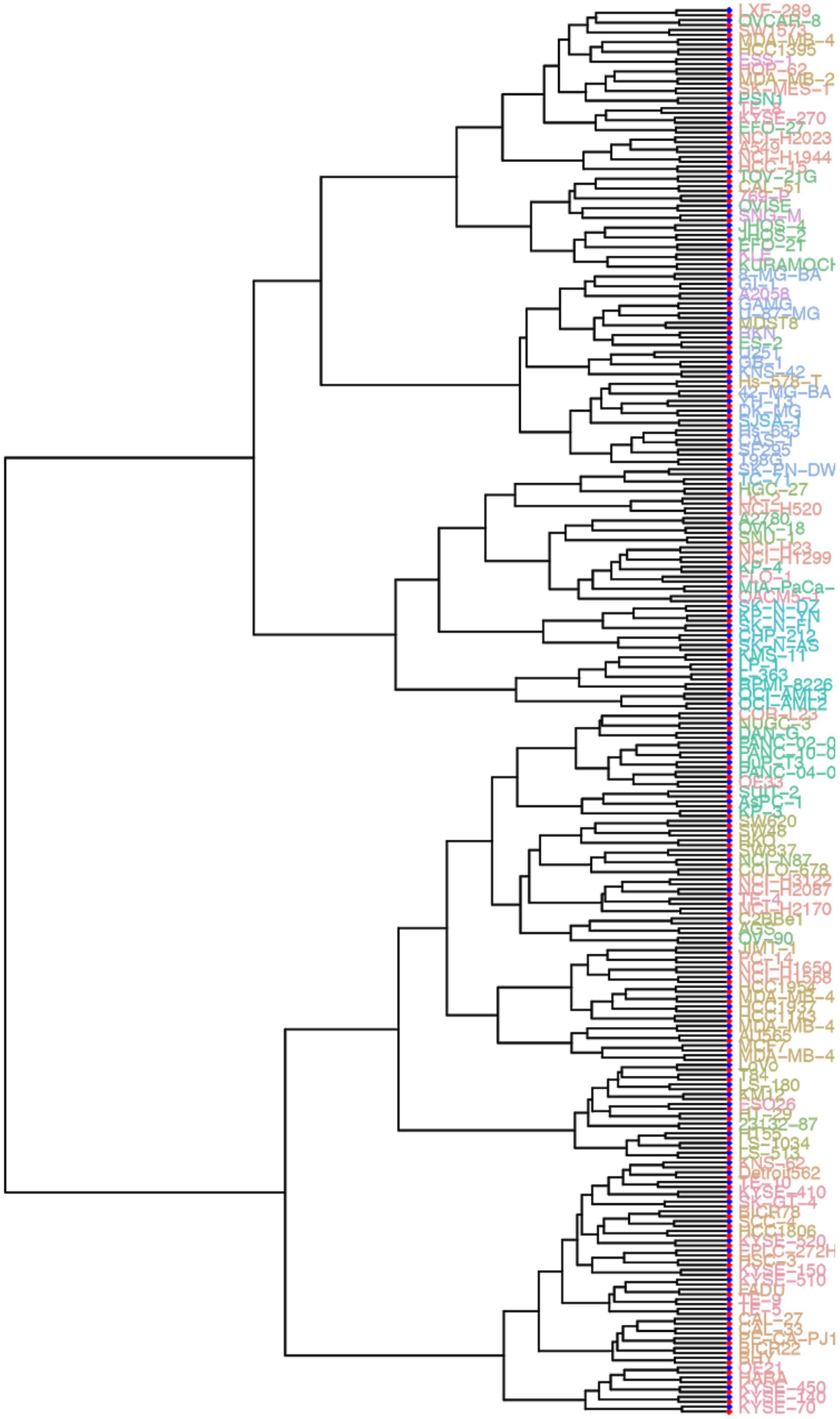

**Supplementary Figure 4**: **Agreement of RNA-seq data sets. (a)** Dendrogram showing the matching of cell lines by institute where the red and blue diamonds and the end of the leaves show the study of origin. The cell line names are colored according to tissue type.

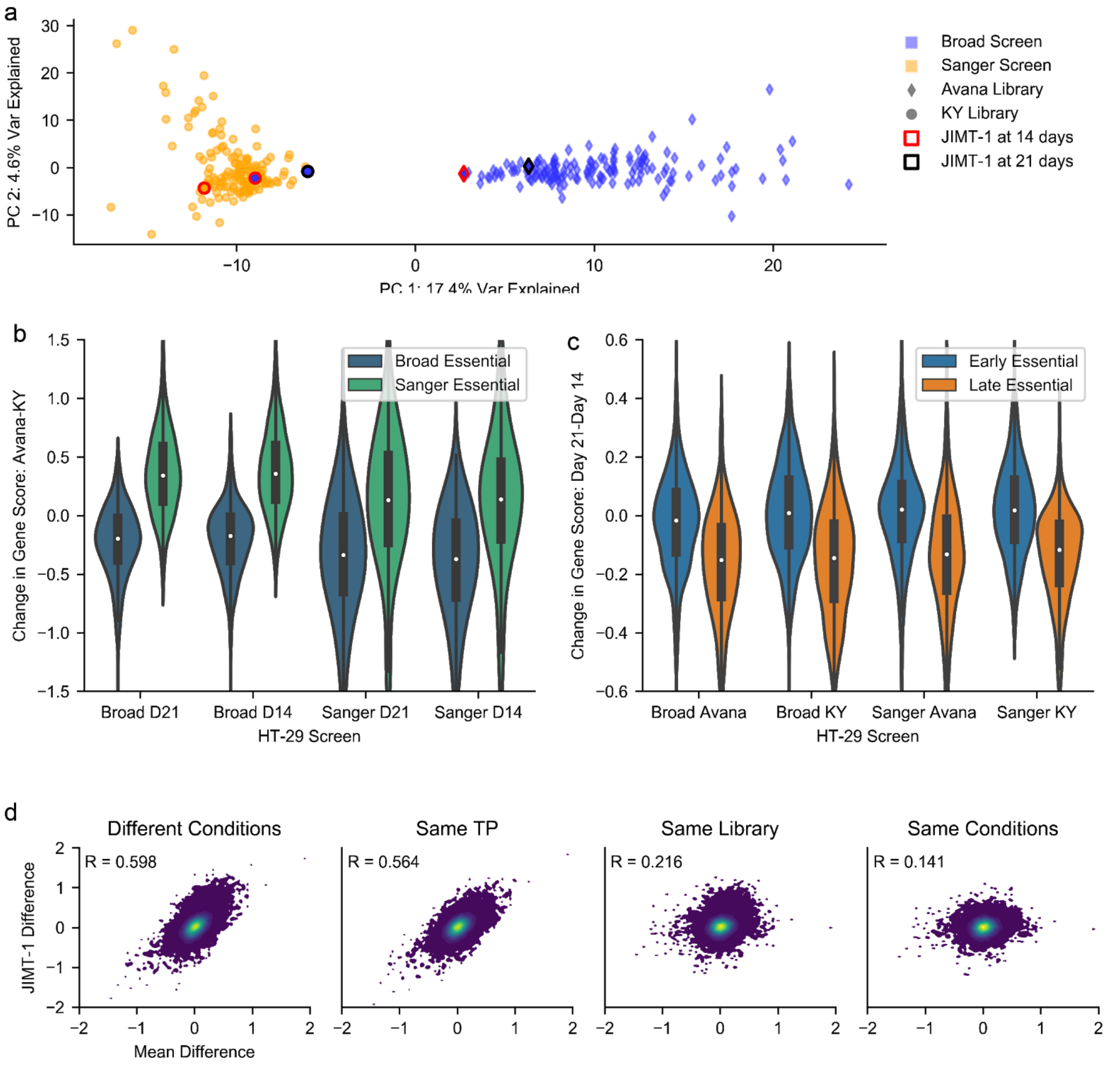


**Supplementary Figure 5**: **Alternate experimental conditions and gene enrichment. (a)** PCA plot of the first two principal components of the concatenated original and replication screens from each institute with JIMT-1 screens highlighted. Axes are scaled to the variance explained by the component. **(b)** Broad- and Sanger-exclusive common dependencies change in mean score when changing libraries under different conditions. X-axis labels indicate timepoint and the institute where the screen was performed. **(c)** Late and early common dependencies change in mean score when changing timepoint under different conditions. X-axis labels indicate library and the institute where the screen was performed. **(d)** The difference in unprocessed gene scores between Sanger screens of HT-29 and the original Broad screen (Broad minus Sanger), beginning with the Sanger's original screen and ending with the Sanger's screen using the Avana library at the 21 day timepoint. Each point is a gene. The horizontal axis is the mean difference of the gene's score between the Broad and Sanger original unprocessed datasets.
